# Is energetics or competition a stronger driver of male smallmouth bass seasonal reproductive timing?

**DOI:** 10.1101/2021.01.30.428956

**Authors:** Robert A.S. Laroche, Kelly L. Weinersmith, Lisa Angeloni, Jeffrey R. Baylis, Steven P. Newman, Scott P. Egan, Daniel D. Wiegmann

**Author notes:** **Corresponding author** Robert A.S. Laroche, G.R.B. W-200I 6100 South Main St, Houston, TX 77005. **LAY SUMMARY** In many animals, larger individuals reproduce earlier in a season. We collected data on reproductive timing in a wild population of bass, and examined whether bigger males breed earlier because they are better competitors, or because they are energetically prepared to breed earlier in the season. Our results suggest that larger males reproduce earlier in a season because they recuperate energy lost in the over-winter period of starvation more rapidly, not because of competitive advantage.

## Abstract

Intraspecific competitive ability is often associated with body size and has been shown to influence reproductive timing in many species. However, energetic constraints provide an alternative explanation for size-related differences of reproductive timing. In temperate fishes that experience a winter starvation period, for instance, a negative allometric relationship between body size and winter energy loss might explain why larger males spawn earlier in a season than smaller males, especially in fishes that exhibit paternal care, which is energetically costly and limits parental foraging opportunities. Male smallmouth bass, *Micropterus dolomieu*, defend nesting territories in which they care for offspring over an extended period. In northern populations, males rely on energy reserves over a winter starvation period and in spring must recoup energy losses before initiating reproduction, making them ideal systems in which to study contributions of competition and energetic allometry on differences of reproductive timing. Here, we harness data on parental male *M. dolomieu* from a 10-year study and show that larger males required fewer degree days-a measure of thermal energy experienced-in spring before they spawned each year and that the time of peak seasonal reproduction in the population was negatively related to the number of degree days accumulated before reproduction started. Furthermore, we found that growth of individual males between seasons better predicted changes in timing of reproduction than changes in size relative to competitors. Together, these results suggest that timing of reproduction in this population is more strongly influenced by energetic constraints than size-based competition amongst males.

## INTRODUCTION

Intraspecific competition has a strong influence on reproductive success in many species (Boeuf 1974; Essington et al. 2000; Wong and Candolin 2005; Boesch et al. 2006; Lane et al. 2009). When competition is between males, body size is often the factor that determines which individuals have a competitive advantage (Alcock 1994; Candolin and Volgt 2001; McElligott et al. 2001; Sih et al. 2002; Newbolt et al. 2016). It comes as no surprise, then, that many studies have found evidence that larger males outcompete smaller males for mates and breed earlier than smaller males, both within and between reproductive seasons (Côte and Hunte 1989; Gibbons 1989; Dufresne et al. 1990; Dickerson et al. 2002; Ciuti and Apollonio 2016).

However, there is an alternative explanation for why bigger individuals of some species breed earlier. In diverse taxa, including fishes, reptiles, insects, amphibians, birds and mammals, larger individuals use less energy proportionate to their mass than smaller individuals (i.e., they show negative allometric relationships with mass specific basal metabolic rate) (Kleiber 1947; Lasiewski and Dawson 1967; Cargnelli and Gross 1997; West et al. 1997; Clarke and Johnston 1999; Hurst and Conover 2003; Nagy 2005; Blanckenhorn et al. 2007; Brodeur et al. 2020). These relationships imply that smaller males will require more time to accumulate energy stores prior to reproduction in species that, say, experience a winter dormant period. Indeed, energetic constraints are known to produce a negative relationship between body size and reproductive timing in a number of species (Tejedo 1992; Danylchuk and Fox 1994; Dobson and Michener 1995; Descamps et al. 2011).

In fishes, where indeterminate growth can lead to substantial variation in size among mature adults, there is an increased opportunity for size to play a role in reproductive timing (Ridgway and Friesen 1992; Knapp et al. 1996; Wiegmann et al. 1997; Dickerson et al. 2005). In some species, within-season variation in timing of reproduction is known to have large fitness consequences (Cargnelli and Gross 1996; Post et al. 1998; Post 2003). Indeed, when males compete for a breeding territory or mates, larger males often have a distinct advantage (Karino 1995; Candolin and Volgt 2001; Stiver and Alonzo 2010). However, a negative allometric relationship to mass specific basal metabolism has also been noted in fishes (Shuter and Post 1990; Cargnelli and Gross 1997; Clarke and Johnston 1999). In north-temperate fishes, where males rely entirely on stored energy reserves throughout winter, such a negative allometric relationship could explain why larger males, which lose proportionately less of their winter energy reserves, spawn earlier in a season than smaller males (Post and Evans 1989; Ridgway et al. 1991; Fullerton et al. 2000). In fishes that exhibit male territoriality and paternal care, both energetics and competitive interactions may contribute to variation in reproductive timing, as is known to occur in other taxa (Langston et al. 1990; Cuadrado and Loman 1999).

Here, we use a 10-year dataset on smallmouth bass, *Micropterus dolomieu*, to tease apart the influence of competition and energetics on reproductive timing. In particular, we follow the reproductive behavior of individual males throughout their lives in a closed population to investigate how male body size, thermal history, and competitive ability impact reproductive timing. There is some evidence that energetics, rather than competition, constrains reproductive timing in *M. dolomieu* (Ridgway et al. 1991; Gillooly and Baylis 1999). However, other aspects of their biology, such as a female preference for larger males and aggressive male-to-male territorial interactions, could also impact reproductive timing (see Ridgway 1988; Ridgway 1989; Baylis et al. 1993; Ongarato and Snucins 1993; Wiegmann et al. 1992; Wiegmann and Baylis 1995). First, we test the hypothesis that larger males reproduce earlier in the spring after the accumulation of fewer degree days, a measure of the thermal energy experienced by an individual. Then, we consider the reproductive timing of the population and test the hypothesis that timing of peak reproduction is negatively related to the number of degree days accumulated before reproduction starts in the spring. Finally, we use comparisons of change in individual male length versus change in competitive status across years to test the hypothesis that an energetic rather than competitive advantage of large body size better predicts changes in within-season reproductive timing.

## MATERIALS AND METHODS

### Natural history

Smallmouth bass, *Micropterus dolomieu*, are native to North America and reproduce seasonally, when water temperatures approach 15 °C (Hubbs and Bailey 1938; Shuter et al. 1980). Males construct nests in the littoral zone into which females deposit eggs. Typically, males mate with only one female and researchers have observed no sneaker behavior (Ridgway 1989; Raffetto et al. 1990; Wiegmann et al. 1992). After spawning, males remain in close proximity to their nest and defend offspring until their fry swim up and disperse, a period that may last several weeks (Hinch and Collins 1991). Parental behavior is costly and leads to a net loss of lean mass, as males rarely feed in this period (Gillooly and Baylis 1999). The energetic cost of parental care in combination with the fact that smaller males end winter with a proportionately larger energy deficit than larger males may allow larger males to spawn earlier within a season than smaller males (Ridgway et al. 1991). In northern populations, smallmouth bass activity ceases when temperatures drop below 10 °C and individuals are largely dormant over winter (Hubbs and Bailey 1938; Munther 1970; Oliver et al. 1979; but see Lyons and Kanehl 2002; Schreer and Cooke 2002).

### Study site

We conducted this study in 1999 and 2001-2009 on Pallette Lake, a 70-ha research lake in the Northern Highlands Fishery Research Area of north-central Wisconsin (46.067° N/89.604° W). The Wisconsin Department of Natural Resources manages this lake, which is closed to fish migration and has a maximum depth of approximately 20 m. Saunders et al. (2002) describe the benthic and limnological characteristics of the lake in detail. Anglers are subject to a mandatory creel census. From 1999-2006, regulations required anglers to return to the lake any male less than 41 cm total length and from 2007 onwards all males under 51 cm had to be returned. From 1999-2006 only 7% of breeding males were larger than 41 cm and from 2007 onward no breeding males were larger than the 51 cm size limit (Table 1).

**Table 1.** Male smallmouth bass and temperature characteristics within and between years.

| Year | Males<br>on<br>Nests | Average<br>Size (cm) | Rank<br>Change | Size<br>Change<br>(cm) | Range<br>of<br>Growth | Winter<br>Length<br>(days) | Mean<br>Temp<br>(°C) | Max<br>Temp<br>(°C) | DD<br>before<br>15 °C |
| --- | --- | --- | --- | --- | --- | --- | --- | --- | --- |
| 1999 | 102 | 31.4 ± 0.7 | NA | NA | NA | NA | 19.3 | 26.7 | 57.4 |
| 2001 | 152 | 29.9 ± 0.6 | NA | NA | NA | 174 | 18.9 | 24.8 | 35.6 |
| 2002 | 221 | 29.8 ± 0.4 | -23.3 ± 1.8 | 5.2 ± 0.2 | 7.5 | 211 | 20.1 | 26.0 | 13.1 |
| 2003 | 234 | 32.3 ± 0.4 | -23.6 ± 1.7 | 5.4 ± 0.2 | 8.3 | 199 | 18.7 | 24.5 | 40.7 |
| 2004 | 284 | 31.6 ± 0.4 | -32.7 ± 1.4 | 4.0 ± 0.1 | 7.0 | 197 | 17.4 | 23.8 | 67.2 |
| 2005 | 216 | 31.9 ± 0.4 | -59 ± 2.1 | 2.6 ± 0.1 | 4.7 | 186 | 19.7 | 27.2 | 20.8 |
| 2006 | 188 | 33.9 ± 0.4 | -43.1 ± 1.8 | 4.8 ± 0.2 | 8.3 | 186 | 19.8 | 25.5 | 27.9 |
| 2007 | 157 | 35.1 ± 0.4 | -39.2 ± 1.6 | 3.7 ± 0.2 | 6.4 | 198 | 18.7 | 25.9 | 16.4 |
| 2008 | 198 | 33.7 ± 0.4 | -20.0 ± 1.3 | 3.0 ± 0.2 | 6.4 | 192 | 18.4 | 23.7 | 32.1 |
| 2009 | 176 | 34.8 ± 0.4 | -23.4 ± 1.4 | 2.5 ± 0.2 | 6.4 | 181 | 17.8 | 23.9 | 77.3 |

Rank change and size (total length) change indicate mean differences ± SE from the previous year. Range of growth indicates the difference between the largest and smallest change in length of an individual from the previous year. Data were not collected in 1998 or 2000 and, hence, these differences were not computed for 1999 or 2001. Degree days (DD) were calculated using the threshold $T_0 = 10\text{ °C}$ . Temperature and Maximum are abbreviated as Temp and Max.

### Nest census

We initiated surveys each year when the water temperature approached 15 °C, typically in mid to late May, and continued until males ceased nest construction (late June to early July) and progeny in most nests had dispersed. Two to three crew members with snorkels searched for nests as they swam transects running from the shoreline out to a depth of about 4 m. This depth was well beyond both the average 1.7 m depth at which nests in the lake were found and the maximum depth found in studies of other Wisconsin lakes (Bozek et al. 2002). Snorkelers placed a numbered tag constructed from a strip of Rite-n-Rain paper tied to a sinker on the edge of each nest and recorded the date, stage of embryos (if present), depth, location and distinctive landmarks that might be useful to relocate a nest when each nest was discovered. They also recorded the developmental stage of any progeny at each subsequent visit to nests, where progressive ontogenetic stages were categorized as eggs, clear fry, grey fry, black-striped fry, black fry or swim-up fry.

### Characteristics of parental males

We captured parental males from their nests with hand nets and recorded their total length (cm). We used uniquely numbered Floy FD-67C anchor tags to track males across years (University of Wisconsin RARC protocol A-48-9700-L00173-2-04-99). We estimated competitors as the number of unique nesting males captured each season (Table 1). We also calculated the average size of these competitors each year. Further, we calculated average growth (cm) and change in competitive ability (i.e., body size, ranked relative to other parental males in a season) for males who bred in consecutive years (Table 1).

### Temperature and degree days

We used temperature data to estimate the thermal energy experienced by parental males prior to reproduction in each year. A thermograph positioned near the shoreline at an approximate depth of 1 m recorded the water temperature (°C) hourly when the lake was not covered by ice. Thermograph malfunctions or other disturbances (e.g., removal of the thermograph from the lake by an unknown person) resulted in some missing temperature records. We estimated these records using regression equations that characterized the relationship between water temperature in Pallette Lake and nearby Sparkling Lake (46.010° N/89.701° W), a North Temperate Lakes Long-Term Ecological Research lake (64 ha; maximum depth 20 m) that has a nearly identical temperature profile as Pallette Lake (Supplement; Supplementary Figure S1; Supplementary Table S1).

Each year we define the growth season as the period between the first day the average temperature climbed above 10 °C and the last day before average temperature dropped below 10 °C (Shuter et al., 1980; Ridgway et al., 1991). We calculated the average and maximum temperature of the growth season for every year. We also calculated winter length, defined as the number of days between growth seasons (Table 1).

We calculated degree days, a measure of the thermal energy experienced by an individual, as the sum of positive differences between the average daily water temperature at 1 m and the threshold T_0_ = 10 °C for each day up to some specified time period (Shuter et al. 1980; Ridgway et al. 1991; Lukas and Orth 1995; Welsh et al. 2017; reviewed by Chezik et al. 2014a, b). In particular, we summed degree days each year from the first date on which the water temperature exceeded 10 °C until the date at which eggs were found in the nest of a male to produce a measure of the seasonal thermal energy experienced by each parental male prior to reproduction (Shuter et al. 1980; Ridgway et al. 1991). In addition, for each year we calculated the number of degree days accumulated by the date that the average water temperature reached 15 °C, the temperature at which reproductive activity is typically initiated. We used this to predict a population-level response time to the rate at which the water temperature increased, where the response time is the interval (days) between the date on which the mean water temperature first reached 15 °C and the median date on which eggs were found in nests (Shuter et al. 1980).

### Statistical analyses

#### Individual behavioral response to temperature

We used linear mixed models to determine the relationship between male total length (log_*e*_ transformed) and the number of degree days (log_*e*_ transformed, threshold T_0_ = 10 °C) that had accumulated up to the date on which a male spawned (i.e., the date a nest was found to contain eggs), where individual was included as a random effect (because 316 individuals bred in more than one season). We fit models with and without year as a covariate and with and without an interaction term between year and male total length and the Akaike information criterion was used to choose between models (Akaike 1973). We restricted the analysis to the first date that a male spawned within a year, if a male spawned more than once within a season (*N* = 127). We excluded males on nests that were discovered after eggs had hatched from the analysis. The total sample included 1,261 observations. If larger males experience less severe winter energy losses, then the slope of the coefficient associated with male length is expected to be negative in each year (Ridgway et al. 1991).

#### Population-level responses to temperature

We used linear regression to predict annual population response times, the time interval between the date on which the mean water temperature reached 15 °C and the median date on which eggs were found in nests, from the number of degree days accumulated before the first 15 °C day each year. If temperature controls individual reproductive readiness, then we expect the slope of this relationship to be negative (Shuter et al. 1980).

#### Energetics versus competition

We used a subset of males (*N* = 265) that spawned in consecutive years to disentangle the potential influences of competition and thermal history on within-season reproductive timing. If the negative allometric association between male body size and winter energy losses controls reproductive timing, then a large increase in male length in a particular year should result in a large decrease in the degree days accumulated before reproduction in the subsequent year. If reproductive timing is controlled by male competitive ability, in contrast, then a male should spawn earlier in a season when confronted with competitors that are relatively small and shifts in competitive status should explain differences of reproductive timing between years. We ranked all males that spawned in each year based on their total lengths, assigning the largest male a rank of zero and smaller males consecutively larger integers (same-sized males received mid-ranks). Notably, this means that a decrease in rank by a male that spawned in two consecutive years corresponds to an increase in size relative to competitors. We used the change of male rank for males that spawned in consecutive years to predict the difference in accumulated degree days before reproduction between years.

We created linear models using either change in rank or change in length (log_*e*_ transformed) between consecutive years to determine the extent to which either factor predicted the respective yearly change in degree days (log_*e*_ transformed) accumulated before individuals reproduced. We also computed the correlation between change in rank and change in length for each year to determine the strength of the association between the two factors (Table 2). Due to the fact that some males appeared in more years than others, it was not possible to include all years in a single analysis with a random effect for individual. Consequentially, we created separate models for each pair of consecutive years to ensure independence of statistics. These models were then compared using the Akaike information criterion (Table 2).

**Table 2.** Linear relationships between change in log_*e*_-transformed male length (ΔL) and competitive status (ΔR) and the change in log_*e*_-transformed degree days (ΔDD) for males breeding in consecutive years.

| Year pair | *N* | | Intercept | Coefficient | Adjusted<br>*R*<sup>2</sup> | *P* | AIC<br>Weight | Correlation<br>($\Delta R$, $\Delta L$) |
| --- | --- | --- | --- | --- | --- | --- | --- | --- |
| 2001-2002 | 46 | $\Delta R$ | -0.443 | 0.004 | -0.008 | 0.420 | 0.09 | *r* = -0.591 |
| | | $\Delta L$ | -0.271 | -1.628 | 0.090 | **0.024** | 0.91 | *P* < 0.001 |
| 2002-2003 | 61 | $\Delta R$ | 0.277 | 0.013 | 0.285 | **< 0.001** | 0.00 | *r* = -0.821 |
| | | $\Delta L$ | 0.611 | -3.680 | 0.465 | **< 0.001** | 1.00 | *P* < 0.001 |
| 2003-2004 | 60 | $\Delta R$ | -0.018 | -0.004 | 0.023 | 0.130 | 0.77 | *r* = -0.867 |
| | | $\Delta L$ | 0.104 | 0.103 | -0.017 | 0.882 | 0.23 | *P* < 0.001 |
| 2004-2005 | 44 | $\Delta R$ | -0.401 | 0.003 | 0.027 | 0.148 | 0.37 | *r* = -0.929 |
| | | $\Delta L$ | -0.424 | -1.482 | 0.050 | 0.078 | 0.63 | *P* < 0.001 |
| 2005-2006 | 49 | $\Delta R$ | -0.101 | 0.003 | 0.051 | 0.064 | 0.34 | *r* = -0.907 |
| | | $\Delta L$ | -0.103 | -0.871 | 0.077 | **0.030** | 0.66 | *P* < 0.001 |
| 2006-2007 | 62 | $\Delta R$ | 0.591 | 0.009 | 0.181 | **< 0.001** | 0.10 | *r* = -0.862 |
| | | $\Delta L$ | 0.548 | -2.779 | 0.238 | **< 0.001** | 0.90 | *P* < 0.001 |
| 2007-2008 | 65 | $\Delta R$ | -0.301 | 0.0002 | -0.016 | 0.931 | 0.50 | *r* = -0.885 |
| | | $\Delta L$ | -0.309 | 0.052 | -0.016 | 0.909 | 0.50 | *P* < 0.001 |
| 2008-2009 | 57 | $\Delta R$ | -0.063 | 0.009 | 0.084 | **0.016** | 0.02 | *r* = -0.891 |
| | | $\Delta L$ | -0.019 | -3.398 | 0.204 | **< 0.001** | 0.98 | *P* < 0.001 |

Degree days were measured up until the point eggs were found in the nest of a male. Bold *P* values indicate *P* < 0.05. The Akaike information criterion is abbreviated as AIC.

## RESULTS

### Characteristics of parental males and seasonal temperature variation

Males were caught over 2,055 nests throughout the course of the study and 847 unique males were tagged. We failed to catch 5.39% of parental males from nests throughout the course of the study. The most males we ever failed to catch in a year was 26 (in 2005). The majority of these instances occurred when offspring died and the nest was abandoned before a male was captured. The number of males on nests ranged from 102 to 284. The total length of males in our data set ranged from 17 cm to 47.5 cm (*N* = 1261) and the seasonal average ranged from 29.8 to 35.1 cm. The mean change in rank and change in size between males that reproduced in consecutive years ranged from - 20.0 to −59 (a negative change in rank indicates an increase in relative size) and 2.5 to 5.4 respectively (Table 1). Temperature patterns also varied year by year. Winter length ranged from 174 days before the 2001 reproductive season to 211 days before the 2002 reproductive season. The mean and maximum temperatures of the growth season ranged from 17.4 to 20.1 °C and 23.7 to 27.2 °C, respectively (Table 1).

### Responses to temperature

#### Individual behavior

In all ten years of the study, a linear relation with negative slope linked male total length (log_*e*_ transformed) to the degree days a male accumulated prior to reproduction (Figure 1). The model that was best supported by the Akaike information criterion had a weight of 0.98, included year and male total length, and no interaction term between these two factors (*N* = 1261, *R*^2^ = 0.63, *P* < 0.0001; Figure 1). The common coefficient for male total length was *b* = −1.27 ± 0.035. The year effect indicates that the y-intercepts differed significantly between seasons (Figure 1).

**Figure 1.**
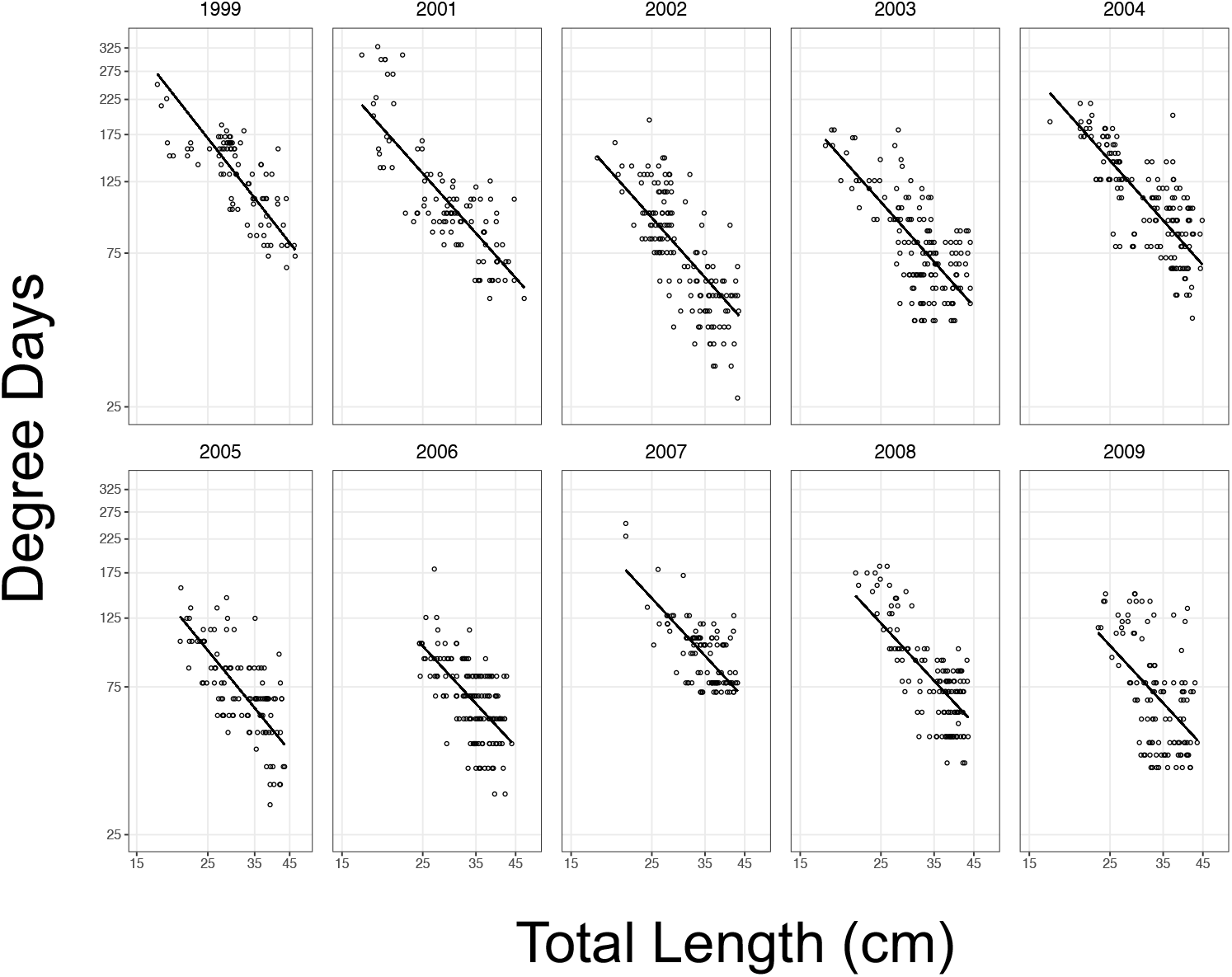
Relationships between parental male length (log_*e*_ transformed) and the degree days accumulated before eggs were observed in nests (log_*e*_ transformed) for 1999, 2001-2009. Data are presented on log-scaled axes.

#### Population-level responses

The dates on which peak reproduction occurred ranged from 23 May in 2001 to 10 June 10 in 2004. The degree days accumulated before the first day that temperatures reached an average of 15 °C varied greatly between years, from 13.1 in 2002 to 77.3 in 2009. In years where more degree days accumulated before the first 15 °C day, the population-level response time was shorter (*N* = 9, *r*^2^ = 0.46, *P* = 0.02; Figure 2). Indeed, in 2009 peak reproduction actually occurred 3 days before the first day that the average temperature reached 15 °C.

**Figure 2.**
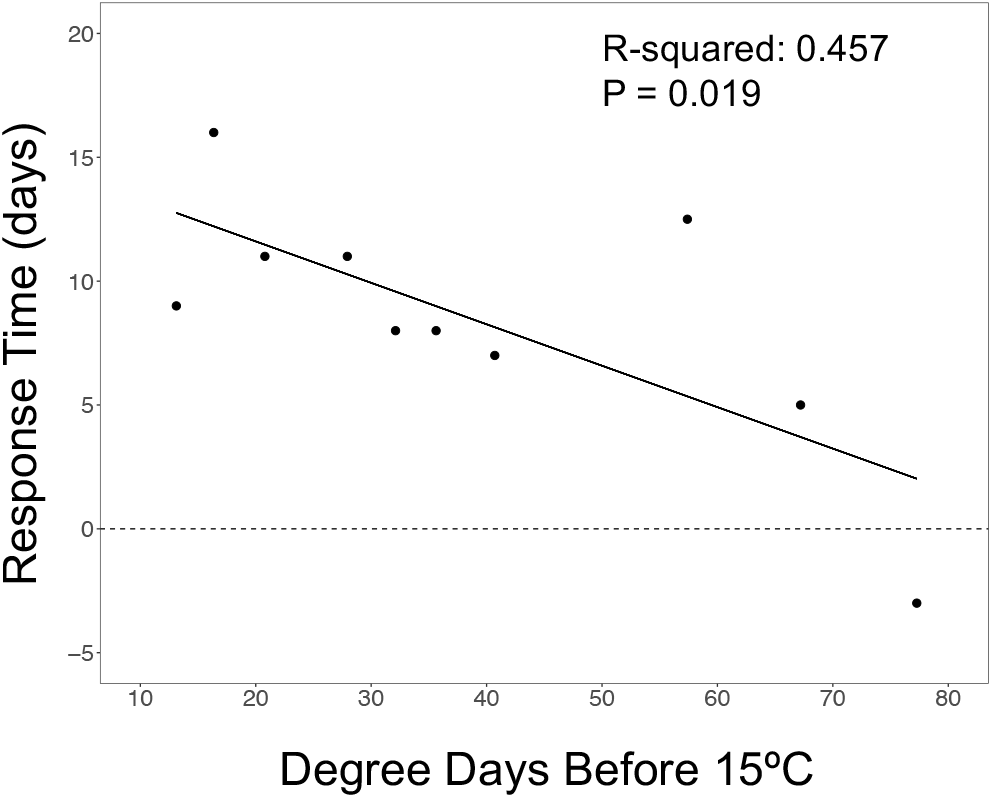
Relationship between the degree days accumulated before the first average daily water temperature of 15 °C (dashed line) and the reproductive response time of the population each year.

#### Energetics versus competition

In 1,009 instances, a male was captured from a nest in consecutive years and for a subset of 444 of these cases—265 unique individuals—the date on which a male spawned (i.e., its nest was discovered with eggs) was known in both years. The difference in male total length between the two years was a significant predictor of the difference in the number of degree days before reproduction in five out of eight (62.5%) of consecutive year pairs, where males that grew more showed larger reductions in degree days before they spawned between the two years (*R*^2^ = 0.08-0.47; Figure 3; Table 2). In comparison, the difference in male rank relative to competitors between the two years was a significant predictor of the change of timing in just three out of the eight (37.5%) of year pairs, where a reduction of body size rank (i.e., increase of competitive status) was associated larger reductions in degree days before they spawned between the two years (*R*^2^ = 0.08-0.29; Table 2). In every year where the relationship between either difference in length or difference in rank was a significant predictor of the difference in the number of degree days before reproduction, models that used difference in length as a predictor received greater weights using the Akaike information criterion (Table 2).

**Figure 3.**
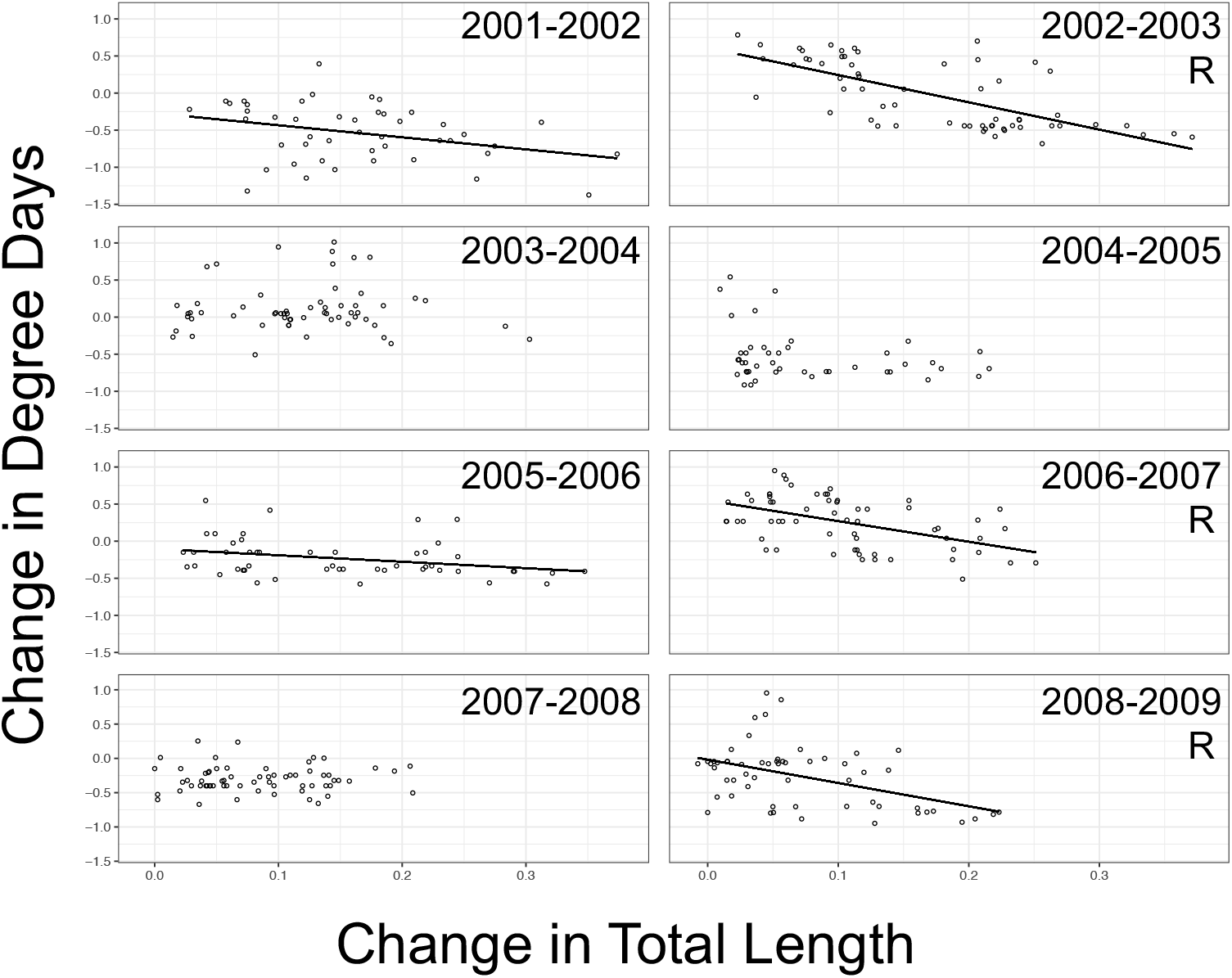
Relationships between change in log_*e*_-transformed total length and change in log_*e*_-transformed degree days before reproduction for males that spawned in consecutive years. Trend lines are included in years where the relationship is significant. The letter R is written under year pairs for which the change of rank was a significant predictor. Data are summarized in Table 2.

Notably, the change of male length between years was a significant predictor in all three year pairs in which the change of male competitive status was a significant predictor. Indeed, in all year pairs—even the two year pairs for which the change in male total length alone predicted the seasonal change of reproductive timing—the two predictors were highly correlated (Table 2). Furthermore, male competitive status appears to have been influential only when the change of male length between years had the strongest associations with changes of reproductive timing (*R*^2^ > 0.2). Together, these results suggest that the ostensible contribution of competitive status on reproductive timing was an artifact of its correlation with male growth between years.

## DISCUSSION

The observed patterns of reproductive timing at both the individual and population level suggest that energetics, rather than competition, explains why larger male *M. dolomieu* spawn earlier in a season than smaller males. Larger males spawned after an accumulation of fewer degree days post winter dormancy than smaller males in all 10 years of the study. At the population level, the delay between the temperature cue that triggers reproduction and the date of peak reproduction was shorter in years where more degree days accumulated prior to the temperature signal. Higher thermal energy experienced by the population appears to drive the distribution of reproduction dates closer to the environmental signal, presumably because males are able to recoup winter energy losses and reach some threshold energetic reserve that is sufficient to carry them through the parental period.

### Energetics over competition

The observed changes of reproductive timing by males that spawned in two consecutive years was better explained by their growth than by changes of their competitive status relative to other males. The change of male total length between years predicted changes of reproductive timing in five of the eight consecutive year pairs we examined, whereas changes of male competitive status—the size-based rank of a male relative to competitors—was a significant predictor of changes in reproductive timing in only three year pairs (Table 2). In the three years where both change in length and change in size-based rank were significant predictors, models using change in length received greater weights based on the Akaike information criterion.

We could identify no social or environmental explanation for why a change in competitive status would be important in some years rather than others (Table 1). The years 2002-2003, 2006-2007 and 2008-2009, where changes of male competitive status appeared to be influential, did not have a larger shift in the number of male competitors than other year pairs, nor did males experience a greater change in status. Especially short winters might in principle relax energetic constraints on reproductive timing and thereby enhance the importance of competition, but none of these year pairs were separated by unusually short winters. Long periods of high temperatures early in the season might have a similar effect and, indeed, 2009 had the highest number of degree days (77.3) accumulated before the average water temperature reached 15 °C. However, this explanation, too, seems unlikely, as 2007 had the second lowest number of degree days (16.4) accumulated before the water temperature reached 15 °C. Lastly, neither the mean nor maximum water temperature, which are known to increase aggressive behavior in fishes, were particularly high in these year pairs (Biro et al. 2010; Colchen et al. 2017; Brandão et al. 2018; Angiulli et al. 2020; Table 1).

The size-based measure of competitive status (i.e., size rank relative to other males) we used presumably also ranks males with regard to female preferences. Hence, female mate preference probably contribute little influence on reproductive timing by males. The female preference for larger males implied in earlier studies by positive correlations between male body size and the number of eggs they acquired seems more likely attributable to a size-related energetic constraint on reproductive timing in females that is similar to that of males (Ridgway et al. 1991; Wiegmann et al. 1992; Baylis et al. 1993).

The effect of growth on changes in reproductive timing was also not observed in all year pairs, but there is a seemingly straightforward explanation, namely that male growth varied considerably between years (Table 1; Figure 3). In year pairs for which no relationship was observed, there were few males that grew especially well and the range of growth was also small, both of which likely impeded the detection of an association (Figure 3). Indeed, the average range of growth in years in which growth was not a significant predictor of changes of reproductive timing was 6.03 cm, compared to 7.38 cm in years where growth was a significant predictor (Table 1).

### Comparisons to previous studies

The control of reproductive timing by individual thermal history is in accord with an earlier study in which parental male removal was not followed by earlier occupancy of nests by smaller males, which would be expected if competition forced smaller males to breed later in the season (Ridgway et al. 1991). There is an additional similarity between our results and those of earlier studies that further supports our hypothesis that reproductive timing in *M. dolomieu* is driven by an underlying metabolic process. Both Ridgway et al. (1991) and Lukas and Orth (1995) examined the association between male total length and degree days accumulated prior to reproduction in Opeongo Lake, Nipissing District, Ontario and North Anna River, Virginia, respectively. In these studies, male length was regressed on degree days (both log transformed) and Lukas and Orth (1995) noted the similarity of the slopes between the two studies, −0.355 and −0.391.

The analysis conducted by Ridgway et al. (1991) included data collected over six years and their analyses indicated that the slope of the relationship did not differ significantly across years. Here, we regressed degree days on male size because we wanted to know how body size influences the number of degree days accumulated by a male before he spawns. But, an equivalent regression analysis with our data (not shown) produced a slope of −0.375, which is remarkably similar to those found in these two earlier studies.

Neuheimer and Taggart (2007) found that a single parameterization of length-at-age as a function of degree days is sufficient to explain growth variation of fish that experience constant or variable water temperatures. The similarity of slopes that characterize the relationship between male body size and accumulated degree days prior to reproduction in geographically separated smallmouth bass systems suggests that reproductive timing is controlled by a conserved metabolic pathway and that a single parameterization based on physiological time—that is, accumulated thermal energy—may explain reproductive timing in diverse systems. Indeed, Shuter and Ridgway (2002) were able to accurately predict *M. dolomieu* start and end reproduction dates based on the regression analysis of Ridgway et al. (1991) in ten water bodies that varied considerably with regard to water temperature and the size distribution of adult males. If competition were the underlying cause of differences in timing of reproduction by males of different sizes, then we would expect the slope to vary both between populations and within populations between years with differences of population density and physical attributes of adult males. The consistent nature of the relationship between male size and degree days before reproduction across the long duration of our study, which encompassed considerable annual variability, provides added confidence that the similarity of slopes between Ridgway et al. (1991) and Lukas and Orth (1995) was not coincidental.

### Applications for management and conservation

For popular game fishes like smallmouth bass, which have a history of management that stretches back nearly 200 years, the identification of broadly consistent behavior between habitats may support current efforts focused on the restoration of native populations (Long et al. 2015). In other high latitude North American fishes, especially those that experience cold winters that impose energetic requirements for individual survival, similar relationships between temperature, size and reproductive timing also likely apply (e.g., Cargnelli and Gross 1997; Hurst and Conover 2003). Changing climate conditions have the potential to greatly disrupt the dynamics of reproductive timing and northern freshwater fish communities are particularly vulnerable to these changes (Visser and Both 2005; Todd et al. 2011; Shuter et al. 2012; Hovel et al. 2017). For example, shorter, less severe winters could relax energetic constraints and permit smaller individuals to reproduce earlier in a season or even at an earlier age, with ample energy reserves, and induce a reversal of control of reproductive timing from one based on energetics to one more dependent on male competitive status. Because the transfer of thermal heat from the environment to fishes accumulates with time, even small changes of temperature over development might cause significant differences in growth trajectories throughout the life of an individual (Neuheimer and Taggart 2007). Further studies that address how energetic constraints and competition influence phenology will be critical to understanding how population dynamics will change as temperatures continue to shift across the globe.

## ACKNOWLEDGMENTS

We thank the University of Wisconsin Trout Lake Biological Station for housing, equipment and other support. Temperature data were provided by Greg Sass and the Wisconsin Department of Natural Resources. For their help in the field we thank S. Bassak, R. Cox, K. Deem, C. Freund, N. Frisch Welch, M. Healey-McNamara, A. Johnson, A. Kautza, K. Kouns, D. Logan, D. Lubin, M. Rice, B. Saetre, V. Schwantes, B. Shefveland, R. Thoni, L. Trayber, J. van Schyndel, J. Weiss, K. Wernert and P. White.

## DATA AVAILABILITY STATEMENT

Data will be deposited in Dryad in accordance with the journal’s data archiving guidelines upon acceptance of this manuscript.

## REFERENCES

Akaike H. 1973. Information theory and an extension of the maximum likelihood principle. Proceedings of the 2nd international symposium on information theory. Second Int Symp Inf Theory.

Alcock J. 1994. Body size and its effect on male-male competition in *Hylaeus alcyoneus* (Hymenoptera: Colletidae). J Insect Behav. doi:10.1007/BF01988901.

Angiulli E, Pagliara V, Cioni C, Frabetti F, Pizzetti F, Alleva E, Toni M. 2020. Increase in environmental temperature affects exploratory behaviour, anxiety and social preference in *Danio rerio*. Sci Rep. doi:10.1038/s41598-020-62331-1.

Baylis JR, Wiegmann DD, Hoff MH. 1993. Alternating life histories of smallmouth bass. Trans Am Fish Soc. doi:10.1577/1548-8659(1993)122<0500:alhosb>2.3.co;2.

Biro PA, Beckmann C, Stamps JA. 2010. Small within-day increases in temperature affects boldness and alters personality in coral reef fish. Proc R Soc B Biol Sci. doi:10.1098/rspb.2009.1346.

Blanckenhorn WU, Fanti J, Reim C. 2007. Size-dependent energy reserves, energy utilization and longevity in the yellow dung fly. Physiol Entomol. doi:10.1111/j.1365-3032.2007.00589.x.

Boesch C, Kohou G, Néné H, Vigilant L. 2006. Male competition and paternity in wild chimpanzees of the Taï forest. Am J Phys Anthropol. doi:10.1002/ajpa.20341.

Bozek MA, Short PH, Edwards CJ, Jennings MJ, Newman SP. 2002. Habitat selection of nesting smallmouth bass *Micropterus dolomieu* in two north temperate lakes. Am Fish Soc Symp.

Brandão ML, Colognesi G, Bolognesi MC, Costa-Ferreira RS, Carvalho TB, Gonçalves-de-Freitas E. 2018. Water temperature affects aggressive interactions in a Neotropical cichlid fish. Neotrop Ichthyol. doi:10.1590/1982-0224-20170081.

Brodeur JC, Vera Candioti J, Damonte MJ, Bahl MF, Poliserpi MB, D’Andrea MF. 2020. Frog somatic indices: Importance of considering allometric scaling, relation with body condition and seasonal variation in the frog *Leptodactylus latrans*. Ecol Indic. doi:10.1016/j.ecolind.2020.106496.

Candolin U, Voigt HR. 2001. Correlation between male size and territory quality: Consequence of male competition or predation susceptibility? Oikos. doi:10.1034/j.1600-0706.2001.950204.x.

Cargnelli LM, Gross MR. 1996. The temporal dimension in fish recruitment: Birth date, body size, and size-dependent survival in a sunfish (bluegill: *Lepomis macrochirus*). Can J Fish Aquat Sci. doi:10.1139/f95-193.

Cargnelli LM, Gross MR. 1997. Notes: Fish energetics: larger individuals emerge from winter in better condition. Trans Am Fish Soc. doi:10.1577/1548-8659(1997)126<0153:nfelie>2.3.co;2.

Chezik KA, Lester NP, Venturelli PA. 2014a. Fish growth and degree-days II: Selecting a base temperature for an among-population study. Can J Fish Aquat Sci. doi:10.1139/cjfas-2013-0615.

Chezik KA, Lester NP, Venturelli PA. 2014b. Fish growth and degree-days I: Selecting a base temperature for a within-population study. Can J Fish Aquat Sci. doi:10.1139/cjfas-2013-0295.

Ciuti S, Apollonio M. 2016. Reproductive timing in a lekking mammal: Male fallow deer getting ready for female estrus. Behav Ecol. doi:10.1093/beheco/arw076.

Clarke A, Johnston NM. 1999. Scaling of metabolic rate with body mass and temperature in teleost fish. J Anim Ecol. doi:10.1046/j.1365-2656.1999.00337.x.

Colchen T, Teletchea F, Fontaine P, Pasquet A. 2017. Temperature modifies activity, inter-individual relationships and group structure in a fish. Curr Zool. doi:10.1093/cz/zow048.

Côte IM, Hunte W. 1989. Male and female mate choice in the redlip blenny: why bigger is better. Anim Behav. doi:10.1016/S0003-3472(89)80067-3.

Cuadrado M, Loman J. 1999. The effects of age and size on reproductive timing in female *Chamaeleo chamaeleon*. J Herpetol. doi:10.2307/1565536.

Danylchuk AJ, Fox MG. 1994. Age and size-dependent variation in the seasonal timing and probability of reproduction among mature female pumpkinseed, *Lepomis gibbosus*. Environ Biol Fishes. doi:10.1007/BF00004929.

Descamps S, Bêty J, Love OP, Gilchrist HG. 2011. Individual optimization of reproduction in a long-lived migratory bird: A test of the condition-dependent model of laying date and clutch size. Funct Ecol. doi:10.1111/j.1365-2435.2010.01824.x.

Dickerson BR, Brinck KW, Willson MF, Bentzen P, Quinn TP. 2005. Relative importance of salmon body size and arrival time at breeding grounds to reproductive success. Ecology. doi:10.1890/03-625.

Dickerson BR, Quinn TP, Willson MF. 2002. Body size, arrival date, and reproductive success of pink salmon, *Oncorhynchus gorbuscha*. Ethol Ecol Evol. doi:10.1080/08927014.2002.9522759.

Dobson FS, Michener GR. 1995. Maternal traits and reproduction in Richardson’s ground squirrels. Ecology. doi:10.2307/1939350.

Dufresne F, Fitzgerald GJ, Lachance S. 1990. Age and size-related differences in reproductive success and reproductive costs in threespine sticklebacks (*Gasterosteus aculeatus*). Behav Ecol. doi:10.1093/beheco/1.2.140.

Essington TE, Quinn TP, Ewert VE. 2000. Intra-and inter-specific competition and the reproductive success of sympatric Pacific salmon. Can J Fish Aquat Sci. doi:10.1139/f99-198.

Fullerton AH, Garvey JE, Wright RA, Stein RA. 2000. Overwinter growth and survival of largemouth bass: interactions among size, food, origin, and winter severity. Trans Am Fish Soc. doi:10.1577/1548-8659(2000)129<0001:ogasol>2.0.co;2.

Gibbons DW. 1989. Seasonal reproductive success of the Moorhen *Gallinula chloropus*: the importance of male weight. Ibis (Lond 1859). doi:10.1111/j.1474-919X.1989.tb02744.x.

Gillooly JF, Baylis JR. 1999. Reproductive success and the energetic cost of parental care in male smallmouth bass. J Fish Biol. doi:10.1006/jfbi.1998.0888.

Hinch SG, Collins NC. 1991. Importance of diurnal and nocturnal nest defense in the energy budget of male smallmouth bass: Insights from direct video observations. Trans Am Fish Soc. doi:10.1080/1548-8659(1991)120[0657:IODANN]2.3.CO;2.

Hovel RA, Carlson SM, Quinn TP. 2017. Climate change alters the reproductive phenology and investment of a lacustrine fish, the three-spine stickleback. Glob Chang Biol. doi:10.1111/gcb.13531.

Hubbs CL, Bailey RM. 1938. The small-mouthed bass. Cranbrook Institute of Science Bulletin 10. 92 pp.

Hurst TP, Conover DO. 2003. Seasonal and interannual variation in the allometry of energy allocation in juvenile striped bass. Ecology. doi:10.1890/02-0562.

Karino K. 1995. Male-male competition and female mate choice through courtship display in the territorial damselfish *Stegastes nigricans*. Ethology. doi:10.1111/j.1439-0310.1995.tb00320.x.

Kleiber M. 1947. Body size and metabolic rate. Physiol Rev. doi:10.1152/physrev.1947.27.4.511.

Knapp RA, Vredenburg VT. 1996. Spawning by California golden trout: characteristics of spawning fish, seasonal and daily timing, redd characteristics, and microhabitat Preferences. Trans Am Fish Soc. doi:10.1577/1548-8659(1996)125<0519:sbcgtc>2.3.co;2.

Lane JE, Boutin S, Gunn MR, Coltman DW. 2009. Sexually selected behaviour: Red squirrel males search for reproductive success. J Anim Ecol. doi:10.1111/j.1365-2656.2008.01502.x.

Langston NE, Freeman S, Rohwer S, Gori D. 1990. The evolution of female body size in red-winged blackbirds: the effects of timing of breeding, social competition, and reproductive energetics. Evolution (N Y). doi:10.1111/j.1558-5646.1990.tb05247.x.

Lasiewski RC, Dawson WR. 1967. A re-examination of the relation between standard metabolic rate and body weight in birds. Condor. doi:10.2307/1366368.

Le Boeuf BJ. 1974. Male-male competition and reproductive success in elephant seals. Integr Comp Biol. doi:10.1093/icb/14.1.163.

Long J, Allen M, Porak W, Suski C. 2015. A historical perspective of black bass management in the United States. Am Fish Soc Symp.

Lukas JA, Orth DJ. 1995. Factors affecting nesting success of smallmouth bass in a regulated Virginia stream. Trans Am Fish Soc. doi:10.1577/1548-8659(1995)124<0726:fansos>2.3.co;2.

Lyons J, Kanehl P. 2002. Seasonal movements of smallmouth bass in streams. Am Fish Soc Symp 31, 149–160.

[dataset] Magnuson, J., S. Carpenter, and E. Stanley. 2020. North Temperate Lakes LTER: High Frequency Meteorological and Dissolved Oxygen Data - Sparkling Lake Raft 1989 - current ver 32. Environmental Data Initiative. https://doi.org/10.6073/pasta/101ae920e5b5a97fadd3cb439440ac3f.

McElligott AG, Gammell MP, Harty HC, Paini DR, Murphy DT, Walsh JT, Hayden TJ. 2001. Sexual size dimorphism in fallow deer (*Dama dama*): Do larger, heavier males gain greater mating success? Behav Ecol Sociobiol. doi:10.1007/s002650000293.

Munther GL. 1970. Movement and distribution of smallmouth bass in the Middle Snake River. Trans Am Fish Soc. doi:10.1577/1548-8659(1970)99<44:madosb>2.0.co;2.

Neuheimer, AB, Taggart, CT. 2007. The growing degree-day and fish size-at-age: the overlooked metric. Can J Fish Aquat Sci 64, 375–385.

Nagy KA. 2005. Field metabolic rate and body size. J Exp Biol. doi:10.1242/jeb.01553.

Newbolt CH, Acker PK, Neuman TJ, Hoffman SI, Ditchkoff SS, Steury TD. 2017. Factors influencing reproductive success in male white-tailed deer. J Wildl Manage. doi:10.1002/jwmg.21191.

Oliver JD, Holeton GF, Chua KE. 1979. Overwinter mortality of fingerling smallmouth bass in relation to size, relative energy stores, and environmental temperature. Trans Am Fish Soc. doi:10.1577/1548-8659(1979)108<130:omofsb>2.0.co;2.

Ongarato RJ, Snucins EJ. 1993. Aggression of guarding male smallmouth bass (*Micropterus dolomieui*) towards potential brood predators near the nest. Can J Zool. doi:10.1139/z93-062.

Post DM. 2003. Individual variation in the timing of ontogenetic niche shifts in largemouth bass. Ecology. doi:10.1890/0012-9658(2003)084[1298:IVITTO]2.0.CO;2.

Post DM, Kitchell JF, Hodgson JR. 1998. Interactions among adult demography, spawning date, growth rate, predation, overwinter mortality, and the recruitment of largemouth bass in a northern lake. Can J Fish Aquat Sci. doi:10.1139/f98-139.

Post JR, Evans DO. 1989. Size-dependent overwinter mortality of young-of-the-year yellow perch (*Perca flavescens*): laboratory, in situ enclosure, and field experiments. Can J Fish Aquat Sci. doi:10.1139/f89-246.

Raffetto NS, Baylis JR, Serns SL. 1990. Complete estimates of reproductive success in a closed population of smallmouth bass (*Micropterus dolomieui*). Ecology. doi:10.2307/1938289.

Ridgway MS. 1988. Developmental stage of offspring and brood defense in smallmouth bass (*Micropterus dolomieui*). Can J Zool. doi:10.1139/z88-248.

Ridgway MS, Shuter BJ, Post EE. 1991. The relative influence of body size and territorial behaviour on nesting asynchrony in male smallmouth bass, *Micropterus dolomieui* (Pisces: Centrarchidae). J Anim Ecol. doi:10.2307/5304.

Ridgway MS. 1989. The parental response to brood size manipulation in smallmouth bass (*Micropterus dolomieui*). Ethology. doi:10.1111/j.1439-0310.1989.tb00728.x.

Ridgway MS, Friesen TG. 1992. Annual variation in parental care in smallmouth bass, *Micropterus dolomieu*. Environ Biol Fishes. doi:10.1007/BF00001890.

Saunders R, Bozek MA, Edwards CJ, Jennings MJ, Newman SP. 2002. Habitat features affecting smallmouth bass *Micropterus dolomieu* nesting success in four northern Wisconsin lakes. Am Fish Soc Symp 31, 123–134.

Schreer JF, Cooke SJ. 2002. Behavioral and physiological responses to smallmouth bass to a dynamic thermal environment. Am Fish Soc Symp 31, 191–203

Shuter BJ, Finstad AG, Helland IP, Zweimüller I, Hölker F. 2012. The role of winter phenology in shaping the ecology of freshwater fish and their sensitivities to climate change. Aquat Sci. doi:10.1007/s00027-012-0274-3.

Shuter BJ, Maclean JA, Fry FEJ, Regier HA. 1980. Stochastic simulation of temperature effects on first-year survival of smallmouth bass. Trans Am Fish Soc. doi:10.1577/1548-8659(1980)109<1:ssoteo>2.0.co;2.

Shuter BJ, Post JR. 1990. Climate, population viability, and the zoogeography of temperate fishes. Trans Am Fish Soc. doi:10.1577/1548-8659(1990)119<0314:cpvatz>2.3.co;2.

Shuter BJ, Ridgway MS. 2002. Bass in time and space: operational definitions of risk. Am Fish Soc Symp 31, 235–249.

Sih A, Lauer M, Krupa JJ. 2002. Path analysis and the relative importance of male-female conflict, female choice and male-male competition in water striders. Anim Behav. doi:10.1006/anbe.2002.2002.

Stiver KA, Alonzo SH. 2010. Large males have a mating advantage in a species of darter with smaller, allopaternal males *Etheostoma olmstedi*. Curr Zool. doi:10.1093/czoolo/56.1.1.

Tejedo M. 1992. Effects of body size and timing of reproduction on reproductive success in female natterjack toads (*Bufo calamita*). J Zool. doi:10.1111/j.1469-7998.1992.tb04454.x.

Todd BD, Scott DE, Pechmann JHK, Whitfield Gibbons J. 2011. Climate change correlates with rapid delays and advancements in reproductive timing in an amphibian community. Proc R Soc B Biol Sci. doi:10.1098/rspb.2010.1768.

Visser ME, Both C. 2005. Shifts in phenology due to global climate change: The need for a yardstick. Proc R Soc B Biol Sci. doi:10.1098/rspb.2005.3356.

Welsh DP, Wiegmann DD, Angeloni LM, Newman SP, Miner JG, Baylis JR. 2017. Condition-dependent reproductive tactics in male smallmouth bass: evidence of an inconsistent birthdate effect on early growth and age at first reproduction. J Zool. doi:10.1111/jzo.12454.

West GB, Brown JH, Enquist BJ. 1997. A general model for the origin of allometric scaling laws in biology. Science (80-). doi:10.1126/science.276.5309.122.

Wiegmann DD, Baylis JR, Hoff MH. 1992. Sexual selection and fitness variation in a population of smallmouth bass, *Micropterus dolomieui* (Pisces: Centrarchidae). Evolution (N Y). doi:10.1111/j.1558-5646.1992.tb01166.x.

Wiegmann DD, Baylis JR. 1995. Male body size and paternal behaviour in smallmouth bass, *Micropterus dolomieui* (Pisces: Centrarchidae). Anim Behav. doi:10.1016/0003-3472(95)80010-7.

Wiegmann DD, Baylis JR, Hoff MH. 1997. Male fitness, body size and timing of reproduction in smallmouth bass, *Micropterus dolomieui*. Ecology. doi:10.2307/2265983.

Wong BBM, Candolin U. 2005. How is female mate choice affected by male competition? Biol Rev Camb Philos Soc. doi:10.1017/S1464793105006809.

