## Supplementary material for "Is energetics or competition a stronger driver of male smallmouth bass seasonal reproductive timing?": Table S1 and Figure S1 (from another version of this preprint: version 1, 2021-02-01)

### SUPPLEMENT

#### Temperature data

Temperatures for Pallette Lake were estimated, when necessary, from data collected at Sparkling Lake, a North Temperate Lakes Long Term Ecological Research site (Magnuson et al. 2020; https://lter.limnology.wisc.edu). The water temperature in the two lakes is highly correlated. For example, the relationship used to predict Pallette Lake water temperature (P) from Sparkling Lake water temperature (S) at 1 m in 1999 was P = -0.1718655 + 1.0351776*S (adjusted $r^2$ = 0.9945; $P$ < 0.0001). All equations used to predict Pallette Lake temperatures from Sparkling Lake temperatures, as well as graphs of combined predicted and measured Pallette Lake temperatures, are provided in Table S1 and Figure S1, respectively.

In 2008, there were some coincident missing temperature data from both lakes. For this year, Sparkling Lake water temperature was iteratively predicted based on the current-day Sparkling Lake air temperature measurements and the water temperature the day before at a depth of 1 m. Sparkling Lake water temperature data were subsequently used to predict Pallette Lake temperatures. Table S1 includes the regression equations used for these calculations. On 43 days, temperature measurements in Pallette Lake changed rapidly between days and deviated substantially (2.5°C or more) from Sparkling Lake temperatures, presumably due to a temporary thermograph malfunction. These anomalies occurred in 2005 and 2007. The mean water temperature in Pallette Lake the day before and the day after these events were averaged to estimate these temperatures. For a few other individual days where water and air temperature were missing for both lakes, the same procedure was used.

**Table S1. Relationships between Pallette Lake (P) and Sparkling Lake (S) water temperatures (ºC) at 1 m depth.**

| Year | Relationship | *Adjusted* $R^2$ | $P$ |
|---|---|---|---|
| 1999 | P = -0.1718655 + 1.0351776*S | 0.9945 | < 0.0001 |
| 2000 | P = 0.6357078 + 0.9897466*S | 0.9931 | < 0.0001 |
| 2001 | P = 0.6655763 + 0.9448224*S | 0.9884 | < 0.0001 |
| 2002 | P = 0.05129394 + 1.00395853*S | 0.9499 | < 0.0001 |
| 2003 | P = 0.05129394 + 1.00395853*S | 0.9499 | < 0.0001 |
| 2004 | P = 0.3855744 + 0.9831321*S | 0.997 | < 0.0001 |
| 2005 | P = -0.9812682 + 1.0459333*S | 0.8878 | < 0.0001 |
| 2006 | P = -0.01003565 + 1.00774548*S | 0.9986 | < 0.0001 |
| 2007 | P = -0.561352 + 1.014938*S | 0.8721 | < 0.0001 |
| 2008 | $S = 0.15162768 + 0.94949871*S_{Prev} + 0.05308003*S_{Air}$ | 0.9977 | < 0.0001 |
| 2008 | P = 0.9820583 + 0.9632688*S | 0.9919 | < 0.0001 |
| 2009 | P = 0.05129394 + 1.00395853*S | 0.9546 | < 0.0001 |

Relationships for 2002, 2003, and 2009 (years for which no Pallette Lake data were available) were determined through a single linear regression of Pallette Lake and Sparkling Lake water temperatures from all the other years combined. In 2008, water temperatures were unavailable on some days for both Pallette and Sparkling Lake. To fill in missing temperature data for Pallette Lake in 2008, missing Sparkling Lake water temperatures were first predicted using air temperatures ($S_{Air}$) and predicted water temperatures from the previous day ($S_{Prev}$).

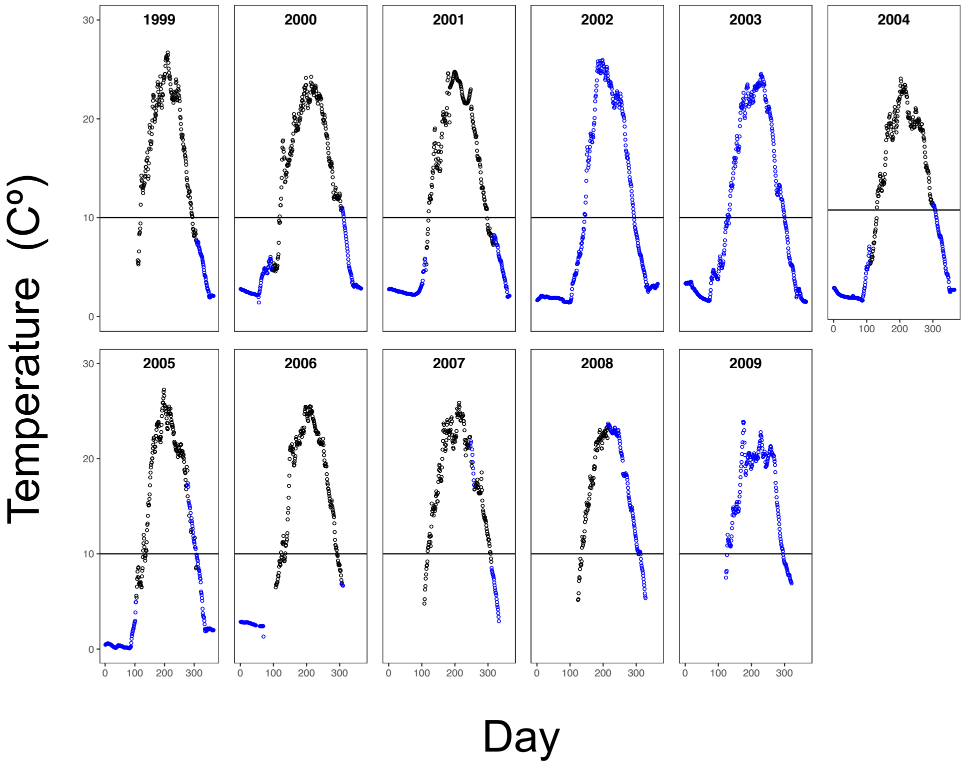


**Figure S1.** Water temperature profiles (1 m depth) for each of the years 1999, 2001-2009. Predicted Pallette Lake temperatures are indicated by blue circles. Predicted temperature data were calculated using the equations in Table S1. Only temperatures above 10 ºC, indicated by a horizontal line on each graph, were relevant to analyses in this study.
